# Chromosome-level reference genome of the European wasp spider *Argiope bruennichi*: a resource for studies on range expansion and evolutionary adaptation

**DOI:** 10.1101/2020.05.21.103564

**Authors:** Monica M. Sheffer, Anica Hoppe, Henrik Krehenwinkel, Gabriele Uhl, Andreas W. Kuss, Lars Jensen, Corinna Jensen, Rosemary G. Gillespie, Katharina J. Hoff, Stefan Prost

**Affiliations:** Zoological Institute and Museum, University of Greifswald, Germany; Institute of Mathematics and Computer Science, University of Greifswald, Germany; Center for Functional Genomics of Microbes, University of Greifswald, Germany; Department of Biogeography, University of Trier, Germany; Interfaculty Institute for Genetics and Functional Genomics, University of Greifswald, Germany; Department of Environmental Science Policy and Management, University of California Berkeley, USA; LOEWE-Centre for Translational Biodiversity Genomics, Germany; South African National Biodiversity Institute, National Zoological Gardens of South Africa, South Africa

**Keywords:** *Argiope bruennichi*, genome assembly, Araneae, spider, PacBio, Hi-C, chromosome-level, whole genome duplication, silk, venom

## Abstract

**Background:** *Argiope bruennichi*, the European wasp spider, has been studied intensively as to sexual selection, chemical communication, and the dynamics of rapid range expansion at a behavioral and genetic level. However, the lack of a reference genome has limited insights into the genetic basis for these phenomena. Therefore, we assembled a high-quality chromosome-level reference genome of the European wasp spider as a tool for more in-depth future studies.

**Findings:** We generated, *de novo*, a 1.67Gb genome assembly of *A. bruennichi* using 21.5X PacBio sequencing, polished with 30X Illumina paired-end sequencing data, and proximity ligation (Hi-C) based scaffolding. This resulted in an N50 scaffold size of 124Mb and an N50 contig size of 288kb. We found 98.4% of the genome to be contained in 13 scaffolds, fitting the expected number of chromosomes (n = 13). Analyses showed the presence of 91.1% of complete arthropod BUSCOs, indicating a high quality of the assembly.

**Conclusions:** We present the first chromosome-level genome assembly in the class Arachnida. With this genomic resource, we open the door for more precise and informative studies on evolution and adaptation in *A. bruennichi*, as well as on several interesting topics in Arachnids, such as the genomic architecture of traits, whole genome duplication and the genomic mechanisms behind silk and venom evolution.

## Data description

### Context

Spider genomes are of great interest, for instance in the context of silk and venom evolution and biomedical and technical applications. Additionally, spiders are fascinating from ecological and evolutionary perspectives. As the most important predators of terrestrial arthropods, they play a key role in terrestrial food webs [1–4]. Spiders are distributed on every continent, except Antarctica, and diverse habitats can be occupied by single species or multiple close relatives [5,6], making them ideal for studies on environmental plasticity, adaptation and speciation. A chromosome-level genome assembly would greatly increase the potential for inference on evolutionary adaptation and modes of speciation [7]. For instance, a well-resolved genome is critical, if evolutionary adaptation happens along genomic islands of differentiation [8–11] or to assess the importance of large genomic rearrangements, such as inversions, in speciation [12–18].

To the best of our knowledge, only eight draft spider genomes have been published to date [19–25], most of which focus on silk and venom genes, while one discusses whole genome duplication [21]. There are three additional, as yet unpublished, spider genome assemblies available on NCBI (National Center for Biotechnology Information) (accession numbers: *Anelosimus studiosus*: GCA_008297655.1; *Latrodectus hesperus*: GCA_000697925.2; *Loxosceles reclusa*: GCA_001188405.1). Spider genomes are considered notoriously difficult to sequence, assemble, and annotate for a number of factors, including their relatively high repeat content, low guanine cytosine (GC) content [19] and due to the fact that they possess some extremely long coding genes in the spidroin gene families [26,27]. Due to these challenges, the completeness of the available spider genomes varies greatly between assemblies (Supplementary Table 1). All of them are incomplete and there is no chromosome-level assembly published for any spider to date. While this does not lessen the conclusions of the above-mentioned studies, a chromosome-level assembly would open doors for more detailed studies on the genomic architecture of gene families, such as silk and venom genes, providing greater understanding of the evolutionary mechanisms driving the diversification of these gene families and genome evolution, in addition to the aforementioned applications in understanding adaptation and speciation.

The European wasp spider, *Argiope bruennichi* (Scopoli, 1772), is an orb-weaving spider in the family Araneidae (Figure 1). Despite the lack of a reference genome, *A. bruennichi* has been the focal species for studies on local adaptation, range expansion, admixture, and biogeography [5,28–30]. These studies have suggested that the range expansion and subsequent local adaptation of *A. bruennichi* to northern Europe was caused by genetic admixture. However, it is not yet known which regions of the genome are admixed, and if these regions are truly responsible for adaptation to colder climates. *A. bruennichi* has also been well studied in the context of dispersal and life history traits [31], as well as sexual selection and chemical communication (e.g. [32–36]). A high-quality reference genome would allow altogether new insights into our understanding of the genetic basis of these phenomena. Considering this background, a chromosome-level reference genome would be highly desirable for the species.

**Figure 1:**
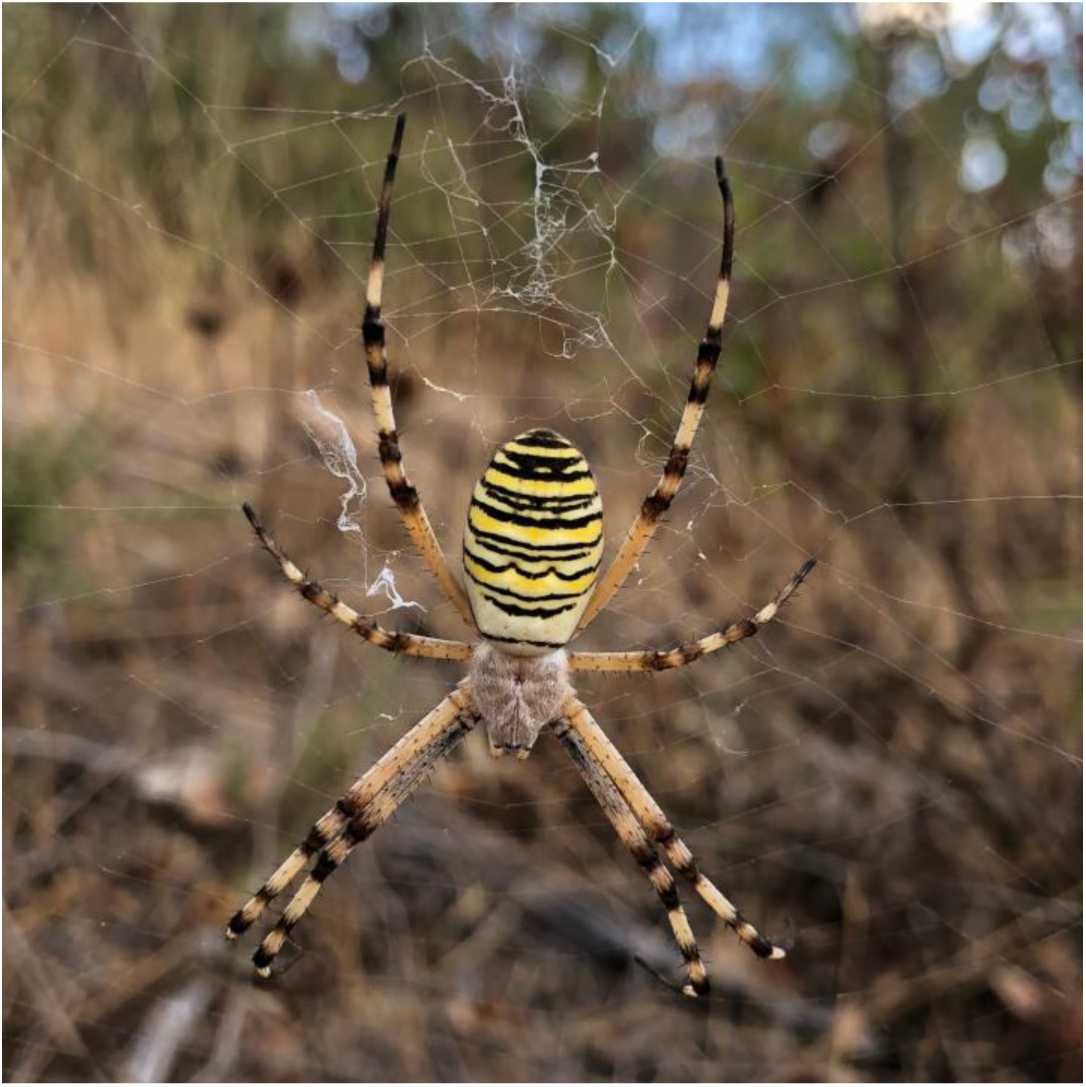
Female *Argiope bruennichi* spider in orb web from Loulé (Faro, Portugal). Photo credit: Monica M. Sheffer

#### Sampling, DNA extraction and sequencing

Adult female *Argiope bruennichi* individuals were collected in the south of Portugal in 2013 and 2019 (Latitude: 37.739 N, Longitude: −7.853 E). As inbred lines of the species do not exist, we selected a population which was previously found to have low heterozygosity in the wild, likely due to naturally high levels of inbreeding [5].

For the baseline assembly, deoxyribonucleic acid (DNA) was extracted from a female collected in 2013 using the ArchivePure blood and tissue kit (5 PRIME, Hamburg, Germany), according to the manufacturer’s protocol. A ribonucleic acid (RNA) digestion step was included using RNAse A solution (7000 U mL^-1^; 5 PRIME). The DNA was stored at −80°C and subsequently sequenced in 2017 at the QB3 Genomics facility at the University of California Berkeley on a Pacific Biosciences Sequel I platform (PacBio, Menlo Park, CA, USA) on 10 cells. The sequencing yielded 21.5X coverage (approximately 36.65 gigabasepairs (Gb), with an estimated genome size of 1.7 Gb).

The specimen collected in 2019 was used to build a proximity-ligation based short-read library (“Hi-C”). Four Hi-C libraries were prepared from a single individual using Dovetail™Hi-C library preparation kit according to the manufacturer’s protocol (Dovetail Genomics, Santa Cruz, CA). The specimen was anesthetized with CO2 before preparation. In brief, the legs were removed from the body and stored in liquid nitrogen, and the leg tissue was disrupted in liquid nitrogen using a mortar and pestle. Chromatin was fixed with formaldehyde, then extracted. Fixed chromatin was digested with DpnII, the 5’ overhangs filled in with biotinylated nucleotides, and the free blunt ends were ligated. After ligation, crosslinks were reversed and the DNA purified to remove proteins. Purified DNA was treated to remove biotin that was not internal to ligated fragments. The DNA was then sheared to ∼350 bp mean fragment size using a Covaris S2 Focused-ultrasonicator. A typical Illumina library preparation protocol followed, with end repair and Illumina adapter ligation. Biotinylated fragments were captured with streptavidin beads before PCR (polymerase chain reaction) amplification (12 cycles), and size selection was performed using SPRI-select beads (Beckman Coulter GmbH, Germany) for a final library size distribution centered around 450 bp. The library was sequenced to approximately 440 million paired end reads on one Flowcell of an Illumina NextSeq 550 with a High Output v2 kit (150 cycles).

#### *De novo* genome assembly

First, we generated a baseline assembly using 21.5X long-read Pacific Biosciences (PacBio) Sequel I sequencing data and the wtdbg2 assembler (v. 2.3) (WTDBG, RRID:SCR_017225) [37]. Next, we polished the assembly by applying three rounds of Pilon (v. 1.23) (Pilon, RRID:SCR_014731) [38] using ∼30X of previously published Illumina paired-end data [5]. This resulted in 13,843 contigs with an N50 of 288.4 kilobase pairs (kb), and an overall assembly size of 1.67 gigabase pairs (Gb). Analysis of Benchmarking Universal Single Copy Orthologs (BUSCO) (v. 3.1.0) scores, using the arthropod data set (BUSCO, RRID:SCR_015008) [39], showed the presence of 90.2% of complete BUSCOs, with 86.4% complete and single-copy BUSCOs, 3.8% complete and duplicated BUSCOs, 3.3% fragmented BUSCOs, and 6.5% missing BUSCOs (Table 1). Next, we scaffolded the contigs using a proximity-ligation based short-read library [40]. Scaffolding using HiRise 2.1.7, a software pipeline designed specifically for using proximity ligation data to scaffold genome assemblies [40], resulted in 13 scaffolds over 1 megabase pairs (Mb) in size, comprising 98.4% of the assembly, with a genome assembly scaffold N50 of 124Mb and BUSCO scores of 91.1% complete genes (Figure 2, Table 1). Genome assembly statistics were calculated using QUAST v. 5.0.2 (QUAST, RRID:SCR_001228) [41] applying default parameters, except --min-contig 0. Previous studies have inferred the chromosome number of *A. bruennichi* to be 13, indicating our genome assembly is full-chromosome level [42,43].

**Table 1:** *Argiope bruennichi* genome assembly completeness.

| Genome assembly statistic | Unscaffolded | Scaffolded |
| --- | --- | --- |
| Assembly size | 1,669,116,561 | 1,670,285,661 |
| AT<sup>a</sup> / GC / N content (%) | 70.7 / 29.3 / 0 | 70.6 / 29.3 / 0.1 |
| **Number of contigs / scaffolds** | 13,843 | 2,231 |
| **Longest contig / scaffold** | 2,039,454 | 143,171,375 |
| **Contig / scaffold N50** | 288,395 | 124,235,998 |
| **Contig / scaffold N90** | 67,231 | 119,022,586 |
| **% repetitive** | 34.66 | 34.64 |
| **BUSCO analysis <sup>b</sup>** |  |  |
| Complete BUSCOs (%) | 90.2 | 91.1 |
| Complete and single-copy BUSCOs (%) | 86.4 | 87.8 |
| Complete and duplicated BUSCOs (%) | 3.8 | 3.3 |
| Fragmented BUSCOs (%) | 3.3 | 2.8 |
| Missing BUSCOs (%) | 6.5 | 6.1 |

Genome assembly statistics were calculated using QUAST v. 5.0.2 (QUAST, RRID:SCR\_001228) [41] using default parameters, except --min-contig 0.
<sup>a</sup> AT: adenine thymine
<sup>b</sup> BUSCO analysis using default parameters against the arthropod dataset

**Figure 2:**
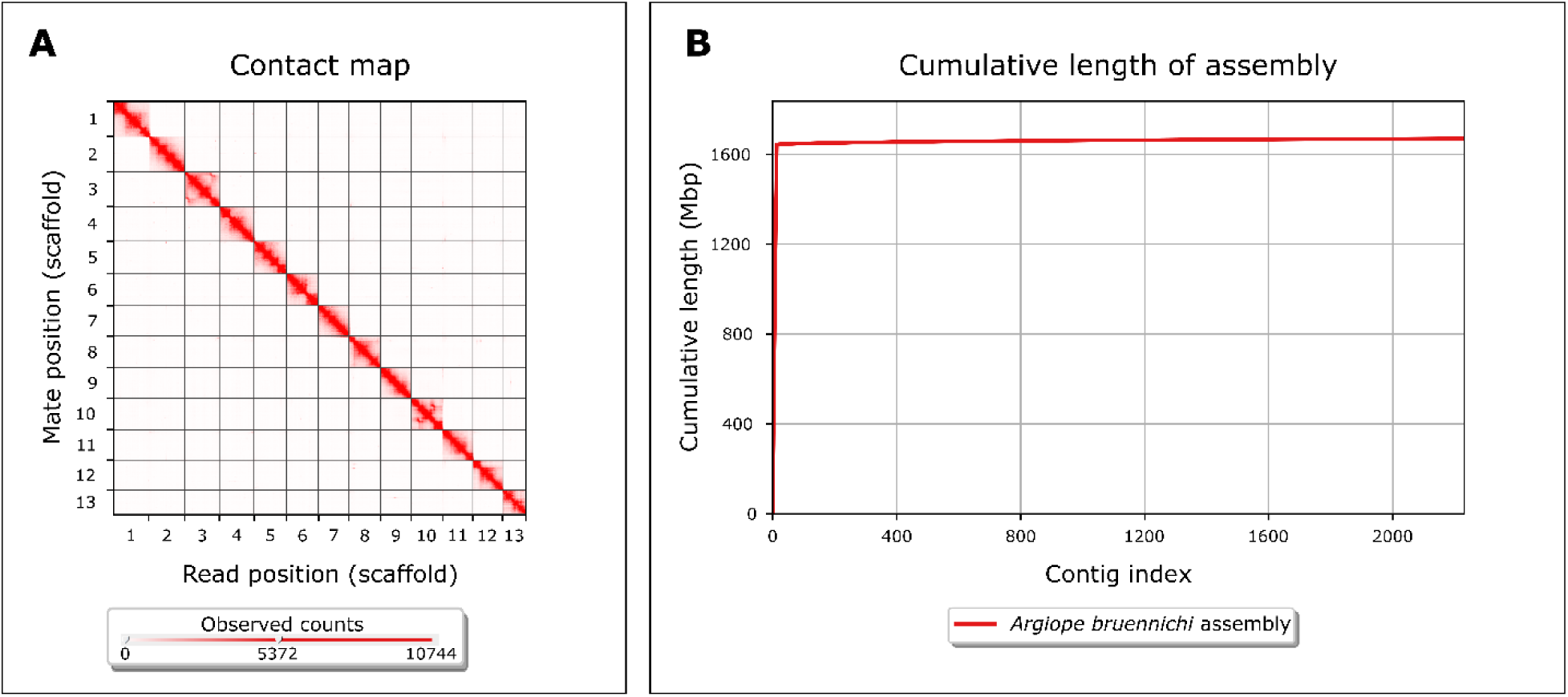
Genome assembly completeness. (A) Contact heatmap of Hi-C scaffolding shows long-range contacts of paired-end Hi-C reads. Gray lines denote scaffold (chromosome) boundaries. Visualized with Juicebox (v. 1.11.08) [66]. (B) Cumulative length of assembly contained within contigs. Note that the vast majority (98.4%) of the genome is contained within very few (13) contigs. Visualized with QUAST v. 5.0.2 [41] using default parameters, except -- min-contig 0.

#### Repeat masking and removal of contaminants

The assembly was repeat-masked using a combination of the *de novo* repeat finder RepeatModeler (v. open-1.0.11) (RepeatModeler, RRID:SCR_015027) [44] and the homology-based repeat finder RepeatMasker (v. open-4.0.9) (RepeatMasker, RRID:SCR_012954) [45]. Repetitive regions accounted for 34.64% of the genome assembly, of which the majority (20.52% of the genome) consisted of unclassified repeats, meaning that they have not been classified in previous studies. The remaining repetitive elements were made up of DNA elements (i.e. transposable elements: 6.27%), long interspersed nuclear elements (LINEs: 1.60%), simple repeats (i.e. duplications of 1-5 bp: 1.58%), long terminal repeat (LTR) elements (0.76%), satellites (0.63%), low complexity repeats (i.e. poly-purine or poly-pyrimidine stretches: 0.42%), and short interspersed nuclear elements (SINEs: 0.08%) (Table 2). BlobTools (v. 1.0) (Blobtools, RRID:SCR_017618) [46] was used to search for contamination, and subsequently mitochondrial sequences and bacterial scaffolds were removed from the assembly. The 14^th^-largest scaffold (Scaffold 839) matched the sequence of a recently-discovered bacterial symbiont of *Argiope bruennichi* [47].

**Table 2:**
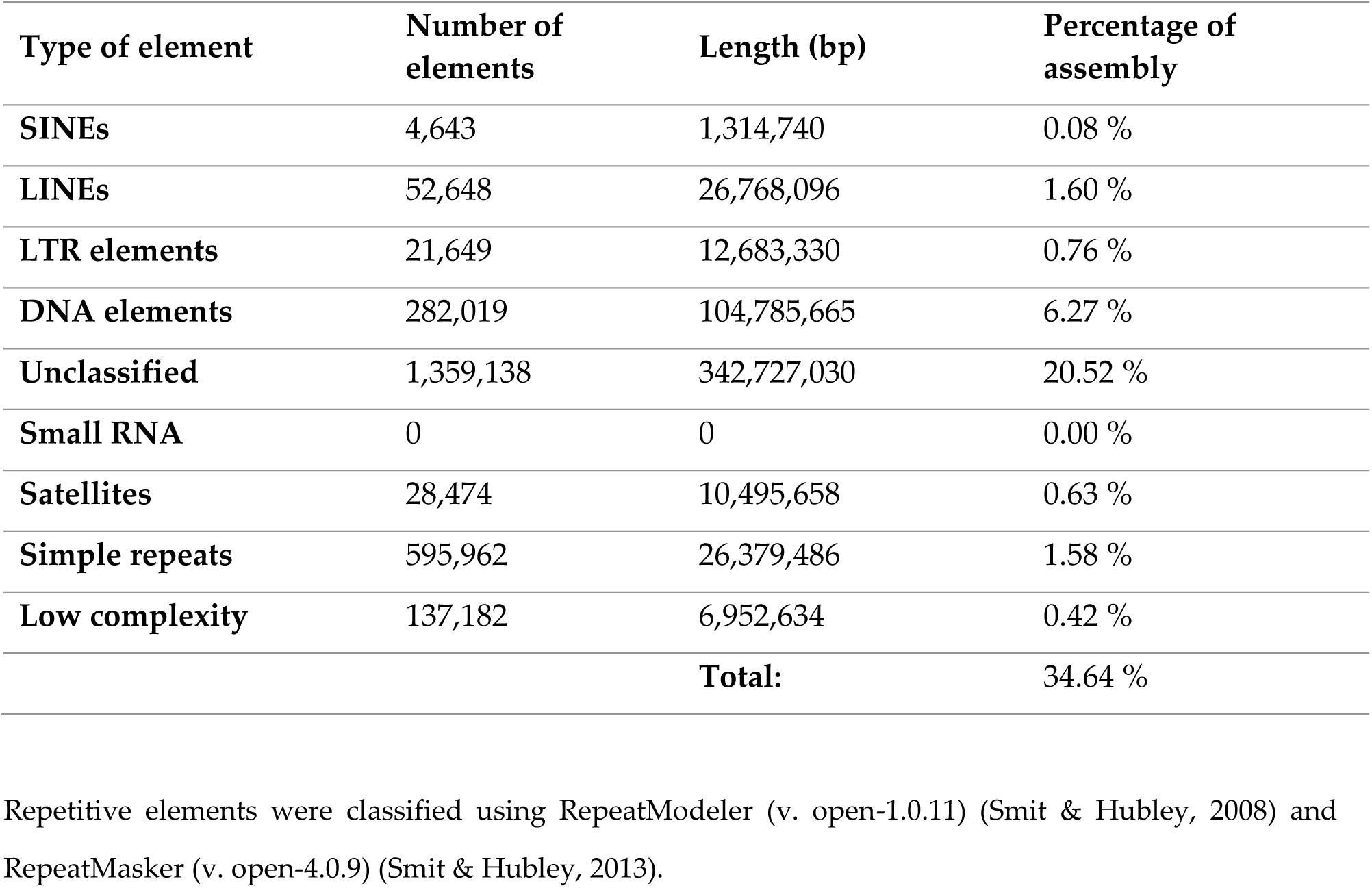
*Argiope bruennichi* repetitive DNA elements.

#### Genome annotation

Raw reads from previously published transcriptome sequencing data [5] were mapped against the repeat-masked assembly using HISAT2 (v. 2.1.0) (HISAT2, RRID:SCR_015530) [48]. After conversion of the resulting SAM file into a BAM file and subsequent sorting using SAMtools (v. 1.7) (SAMTOOLS, RRID:SCR_002105) [49], the sorted BAM file was converted to intron-hints for AUGUSTUS (v. 3.3.2) (Augustus, RRID:SCR_008417) [50] using AUGUSTUS scripts. AUGUSTUS was run on the soft-masked genome with the *Parasteatoda* parameter set. The resulting gff file containing predicted genes was converted into a gtf file using the AUGUSTUS script gtf2gff.pl. Additional AUGUSTUS scripts (getAnnoFastaFromJoinGenes.py and fix_in_frame_stop_codon_genes.py) were used to find and replace predicted genes containing in-frame stop codons with newly predicted genes. The resulting gtf file containing 23,270 predicted genes was converted to gff3 format using gtf2gff.pl and protein sequences of predicted genes were extracted with getAnnoFastaFromJoinGenes.py. Finally, functional annotation was performed using InterProScan (v. 5.39-77.0) (InterProScan, RRID:SCR_005829) [51,52] (Table 3).

**Table 3:** *Argiope bruennichi* genome annotation statistics.

| **Genome Annotation Statistic** | **Value** |
| --- | --- |
| **Number of protein coding genes** | 23,270 |
| **Functionally annotated genes (%)** | 81.0 |
| **Average exon length (bp)** | 200 |
| **Average intron length (bp)** | 4,035 |
| **BUSCO analysis <sup>a</sup>** |  |
| Complete BUSCOs (%) | 89.3 |
| Complete and single-copy BUSCOs (%) | 76.7 |
| Complete and duplicated BUSCOs (%) | 12.6 |
| Fragmented BUSCOs (%) | 7.0 |
| Missing BUSCOs (%) | 3.7 |

<sup>a</sup> BUSCO analysis using default parameters against the arthropod dataset

#### Comparative genomic analysis of repeat content

High repetitiveness is characteristic of spider genomes [19]. In order to compare the repeat content of *A. bruennichi* with that of other spiders, we downloaded the genome assemblies of several other spider species from NCBI and the DNA Data Bank of Japan (DDBJ) (accession numbers in Table 4), then treated them in the same manner as the *A. bruennichi* genome, masking the repeats using RepeatModeler (v. open-1.0.11) [44] and RepeatMasker (v. open-4.0.9) [45]. *Acanthoscurria geniculata* was excluded from this analysis due to the very large and relatively poorly assembled genome. The *A. bruennichi* genome has a slightly lower percentage of repetitive element content (34.64%) compared to most other spiders (Table 4). Some species, such as *Loxosceles reclusa, Trichonephila clavipes* (formerly *Nephila clavipes*), *Anelosimus studiosus* and *Parasteatoda tepidariorum*, have similar repetitive content (36.51%, 36.61%, 35.98% and 36.79% respectively); other species have much higher repetitive content, such as *Araneus ventricosus, Dysdera silvatica, Stegodyphus dumicola, Stegodyphus mimosarum* and *Pardosa pseudoannulata* (55.96%, 60.03%, 58.98%, 56.91% and 48.61% respectively). Only *Latrodectus hesperus* has lower repetitive content (20.97%). The classification and relative percentage of these repeats can be found in Supplementary Table 2 and Supplementary Figure 1.

**Table 4:** Total repetitive content in the genomes of spiders.

| Species | % repetitive | Accession number [reference] |
| --- | --- | --- |
| *Argiope bruennichi* | 34.64 | -- |
| *Araneus ventricosus* | 55.96 | BGPR01000001-BGPR01300721 <sup>a</sup> [22] |
| *Trichonephila clavipes* | 36.61 | GCA_002102615.1 <sup>b</sup> [20] |
| *Dysdera silvatica* | 60.03 | GCA_006491805.1 <sup>b</sup> [25] |
| *Stegodyphus dumicola* | 58.98 | GCA_010614865.1 <sup>b</sup> [24] |
| *Stegodyphus mimosarum* | 56.91 | GCA_000611955.2 <sup>b</sup> [19] |
| *Pardosa pseudoannulata* | 48.61 | GCA_008065355.1 <sup>b</sup> [23] |
| *Loxosceles reclusa* | 36.51 | GCA_001188405.1 <sup>b</sup> [unpublished] |
| *Anelosimus studiosus* | 35.98 | GCA_008297655.1 <sup>b</sup> [unpublished] |
| *Latrodectus hesperus* | 20.97 | GCA_000697925.2 <sup>b</sup> [unpublished] |
| *Parasteatoda tepidariorum* | 36.79 | GCA_000365465.3 <sup>b</sup> [21] |

Repetitive elements were classified using RepeatModeler (v. open-1.0.11) (Smit & Hubley, 2008) and RepeatMasker (v. open-4.0.9) (Smit & Hubley, 2013).
<sup>a</sup> DNA Data Bank of Japan (DDBJ)
<sup>b</sup> National Center for Biotechnology Information (NCBI)

#### Genome architecture of Hox, spidroin and venom genes

Previous studies on spider genomes have focused on whole genome duplication, silk gene evolution, and venom gene evolution [19–23]. Therefore, to place the *A. bruennichi* genome into the same context, we manually curated three gene sets from publicly available protein sequences: Hox, spidroin (silk), and venom genes. Because Hox genes are highly conserved across taxa [53], we chose the most complete sequences for the ten arthropod Hox gene classes from spiders without regard to the relatedness of the species to *A. bruennichi* (Supplementary Table 3). In contrast to Hox genes, spidroin and venom genes are highly polymorphic and species-specific [54–57]. For the spidroin gene set, we downloaded protein sequences of the seven spidroin gene classes exclusively from five species of the genus *Argiope* (Supplementary Table 4). Venom genes are best studied in spiders that are medically significant to humans, which are very distant relatives to *A. bruennichi* [51–54]. To allow comparison, we focused on venom gene sequences available for araneid spiders (two species, Supplementary Table 5); however, the function and classification of these genes is poorly understood. With these three gene sets (Hox, spidroin, and venom), we performed a TBLASTN search against our genome assembly (v. 2.10.0+) (TBLASTN, RRID:SCR_011822) [62,63]. We recorded the genomic position of the best matches and compared them with the AUGUSTUS gene predictions for those locations. We employed a conservative E-value cutoff of less than 1.00E-20 and only included results with an identity greater than 60%. The manually curated FASTA files of each gene set used for the TBLASTN search are available in Supplementary Files 1-3. A table of the matches with accession numbers for each gene set is available in Supplementary Tables 3-5.

### Whole genome duplication

In 2017, Schwager *et al.* asserted that a whole genome duplication (WGD) event occurred in the ancestor of scorpions and spiders, as evidenced by a high number of duplicated genes, including two clusters of Hox genes in *Parasteatoda tepidariorum* and the bark scorpion *Centruroides sculpturatus* [21]. In their study, they found one nearly-complete cluster of Hox genes on a single scaffold, lacking the *fushi tarazu (ftz)* gene, which they argued may be the case for this cluster in all spiders. The second set of Hox genes was distributed across two scaffolds, which the authors attributed to incompleteness of the assembly due to patchy sequencing coverage [21]. For consistency, we will use the same nomenclature for Hox genes as used in [21] (*Abdominal-B: AbdB, Abdominal-A: AbdA, Ultrabithorax: Ubx, Antennapedia: Antp, fushi tarazu: ftz, sex combs reduced: scr, Deformed: Dfd, Hox3, proboscipedia: pb, labial: lab*). Corresponding with the results from *P. tepidariorum*, we found two clusters of Hox genes, with no evidence of tandem duplication. The two clusters occurred on two chromosomes (Chromosome 9 and Chromosome 6). In these locations, InterProScan generally annotated the genes as Hox genes but did not identify the specific type. On Chromosome 9, the Hox genes were in reverse collinear order, with no overlapping regions. Because it is complete, we will refer to this cluster as “Cluster A.” On Chromosome 6, (“Cluster B”) the genes were out of collinear order, with the position of *AbdA* and *Ubx* switched, and the coordinates for *Dfd, Hox3* and *pb* from the blast search overlapping (Figure 3A). The hits for *Antp* and *ftz* in Cluster B fell onto a single predicted gene in the annotation. Thus, it is unclear if *A. bruennichi* lacks one copy of *ftz*, as in *P. tepidariorum*, or if the annotation incorrectly fused the two genes in this cluster. In the study by Schwager *et al.*, 2017 [21], low sequencing coverage of Cluster B downstream of *Dfd* limited their inference. In our genome assembly, by mapping the PacBio reads against the final assembly, we calculated that we have an average of more than 12X coverage across the length of both clusters, suggesting that Cluster B is not out of order due to problems arising from low coverage. It is possible that Hox Cluster B in spiders has changed or lost functionality following the ancestral WGD event.

**Figure 3:**
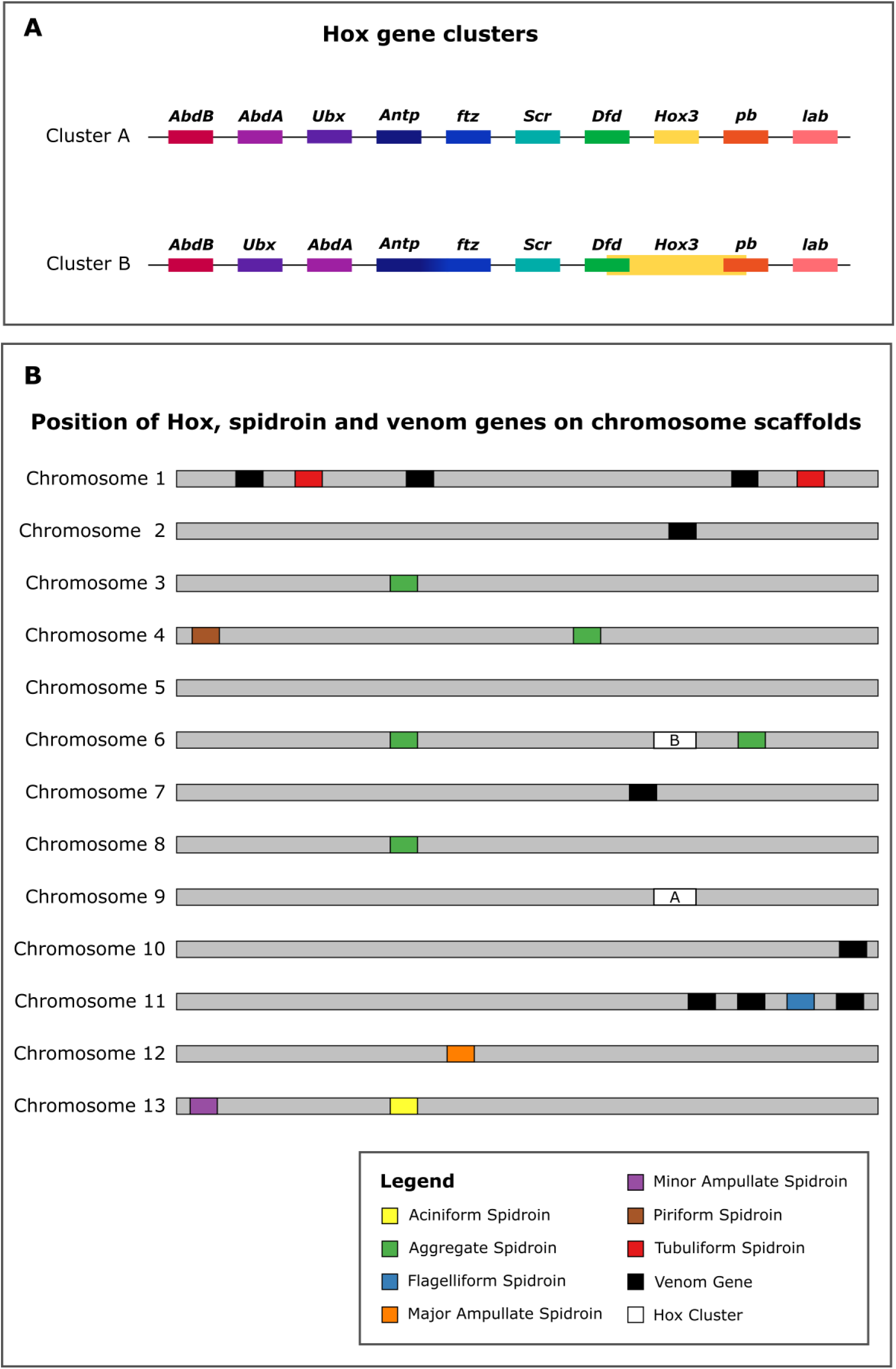
Schematic representation of location of gene families on the 13 chromosomes. (A) Hox gene clusters. Genes connected by a black line occur on the same scaffold. Cluster A occurs on Chromosome 9, and Cluster B occurs on Chromosome 6. The presence of two Hox gene clusters on two chromosomes validates the previous finding of whole genome duplication [21]. (B) Position of Hox, spidroin and venom genes on chromosome scaffolds. The light grey bars represent chromosomes, the colored rectangles represent the seven different spidroin gene families, the black rectangles represent venom genes, and the white rectangles represent Hox gene clusters.

### Spidroin genes

There are seven classes of silk produced by araneomorph spiders, each with one or more unique uses; it is important to note that the uses of these silk types are best understood for spiders in the family Araneidae, and the number and uses of silk types can vary widely between families [20,22,64,65]. The classes of silk are major ampullate (*MaSp*) minor ampullate (*MiSp*), piriform (*PiSp*), aggregate (*AgSp*), aciniform (*AcSp*) tubuliform (also referred to as cylindrical) (*TuSp*) and flagelliform (*Flag*). In *A. bruennichi*, spidroin genes occur on eight out of the thirteen chromosome scaffolds (Chromosomes 1, 3, 4, 6, 8, 11, 12 and 13) (Figure 3B). There were no hits on smaller scaffolds. In the majority of cases, all blast matches for a single spidroin type occurred on a single chromosome; the only exception was for *AgSp*, which had hits on four different chromosomes. However, these were not all annotated as spidroins; on Chromosome 6 there were multiple hits which were annotated as spidroins, while on Chromosome 4 the hit was annotated as tropoelastin, on Chromosome 3 the hit was annotated as a chitin binding domain, and on Chromosome 8 the hit was annotated as a serine protease. All hits for *TuSp* occurred on Chromosome 1, but there were two physically separated clusters on the chromosome. There are more sequences available for *MaSp* than any of the other spidroin types in the genus *Argiope*, which allowed us to find matches for several unique *MaSp* genes in the *A. bruennichi* assembly. These occur in a small region of Chromosome 12, in close proximity to one another, suggesting that the spidroin genes in this group may have diversified via tandem duplication.

### Venom genes

We found high identity matches for venom toxins on five of the chromosome scaffolds (Chromosomes 1, 2, 7, 10 and 11) (Figure 3B), but the majority of hits were on Chromosome 1. Babb *et al.* 2017 conducted a study on silk genes in *Trichonephila clavipes* (formerly *Nephila clavipes*), in which they found a novel flagelliform-type gene (FLAG-b) which was expressed most highly in the venom glands, not the flagelliform silk glands. This added to previous findings in the *Stegodyphus mimosarum* genome, where spidroin-like proteins in the venom glands are found [19]. Interestingly, in the *A. bruennichi* genome assembly, there are several venom genes on Chromosome 11 in close proximity to flagelliform spidroin genes.

### Conclusion

We have assembled and annotated the first chromosome-level genome for an arachnid. The assembly approach of combining long read, short read, and proximity ligation data overcame the challenges of assembling arachnid genomes, namely genome size, high repetitiveness, and low GC content. In our study, we made a preliminary analysis of the location of certain gene families of interest in the context of spider genomics, which hinted at several interesting directions for future studies on the evolution of silk and venom genes. Furthermore, because this species has undergone a recent and rapid range expansion, the well-resolved genome assembly will be useful for studies on the genomic underpinnings of range expansion and evolutionary adaptation to novel climates.

## Availability of supporting data

The final genome assembly and raw data from the PacBio and Hi-C libraries, as well as the annotation, have been deposited at NCBI under BioProject PRJNA629526 and will be available upon publication. A publicly accessible genome browser hub with the annotation and raw transcriptome and PacBio read coverage can be found on the UCSC Genome Browser server (hub name “Wasp spider hub”).

## Availability of source code and requirements

All data required to replicate this work are available on NCBI and in the supplementary files.

## Declarations

## List of abbreviations

*Abd-A*: *Abdominal-A*;
*Abd-B*: *Abdominal-B*;
*AcSp*: aciniform spidroin;
*AgSp*: aggregate spidroin;
*Antp*: *Antennapedia;*
AT: adenine thymine;
bp: basepairs;
BUSCO: Benchmarking Universal Single Copy Orthologs;
DDBJ: DNA Data Bank of Japan;
*Dfd*: *Deformed;*
DNA: deoxyribonucleic acid;
*Flag*: flagelliform spidroin;
*ftz*: *fushi tarazu;*
Gb: gigabase pairs;
GC: guanine cytosine;
kb: kilobase pairs;
*lab*: *labial;*
LINE: long interspersed nuclear element;
LTR: long terminal repeat;
*MaSp*: major ampullate spidroin;
Mb: megabase pairs;
*MiSp*: minor ampullate spidroin;
NCBI: National Center for Biotechnology Information;
PacBio: Pacific Biosciences;
*pb*: *proboscipedia;*
PCR: polymerase chain reaction;
*PiSp*: piriform spidroin;
RNA: ribonucleic acid;
*scr*: *sex combs reduced;*
SINE: short interspersed nuclear element;
*TuSp*: tubuliform spidroin;
*Ubx*: *Ultrabithorax;*
WGD: whole genome duplication

## Consent for publication

Not applicable.

## Authors’ contributions

MMS, HK, GU, and SP conceived of the study; MMS, HK, and GU collected the spiders. HK extracted DNA for the PacBio sequencing; MMS prepared and submitted the DNA for PacBio sequencing, with input and infrastructure provided by RGG. MMS and CJ constructed and sequenced the Hi-C library, with input and infrastructure provided by LJ and AK. MMS, AH and SP performed the genome assembly, and AH and KJH performed the genome annotation with input and infrastructure provided by MMS and SP. AH analyzed the repeat content of other spider genomes; MMS performed the analysis of whole genome duplication, spidroin genes, and venom genes. MMS, AH, KJH and SP wrote the first draft of the manuscript. All authors read and approved the final manuscript.

## Acknowledgements

We would like to thank the California Academy of Sciences for allowing us access to their computing resources for the genome assembly, and to Dovetail Genomics for their support in troubleshooting the Hi-C kit and running HiRise. MMS thanks José Cerca for helpful ideas and discussions about the silk and venom gene analysis.

