## Supplementary figures and images for "Chromosome-level reference genome of the European wasp spider *Argiope bruennichi*: a resource for studies on range expansion and evolutionary adaptation"

### Supplementary Figure 1

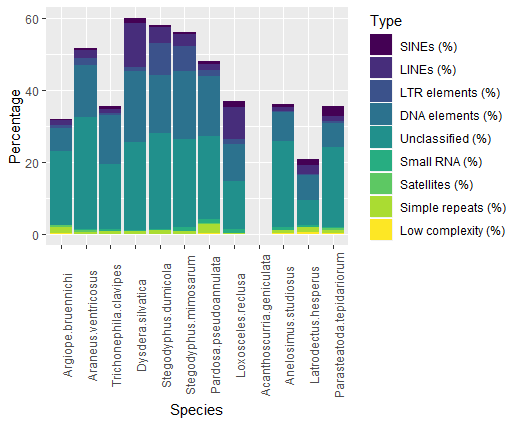
